# Transient dynamics response analysis of the limb under impact loading

**DOI:** 10.1101/2024.08.22.609134

**Authors:** Mengjun Song, Jinggong Wei, Liping Zhang

## Abstract

The limb vibrations are initiated at paw-strike in animal’s normal movement. The short ground contact duration suggests that there exists the transient dynamic response of the impact between the leg and ground, which is a high nonlinear problem and not well understood. The purpose of this study is to investigate the transient dynamic response of the lower limb of the felid during the moving. A kinematic model of the musculoskeletal system of a small felid is constructed from the anatomical measurement data. The elastic moduli are measured and calculated for different parts of the lower limb by a Nano-indentation technique. On the basis of the measured material parameters, the substructure technique is employed to numerically solve the contact-impact behavior of the lower limb. The high speed imaging system and a designed electronic detection system are employed in the experiment to testify the numerical results, which demonstrate that the initial impact has an important influence on the performance of the lower limb during the movement especially the high speed movement, and the paw-pad has good damping effects during the impact. The multiple impacts exist between limbs and the soil, which may provide the feedback energy for the felid’s high-speed running, though it may also increase the risk of the stress fracture of the limbs.

## Introduction

When a quadruped walks or gallops, the lower extremities of its forelimbs and hind limbs alternately contact the ground. Especially during galloping, contact-impact as a transient response process exists between the lower extremities and the ground ^[1, 2]^. Therefore, typical time histories of the initial contact event during the impact are usually very short. The impact force, caused by inertial change of the lower extremity over the gait cycle, reaches a larger value than the body weight^[3]^, and generates the dynamic response wave^[4]^ transmitted along the skeletons and muscles^[5,6,7]^. However, the impact shock during contact-impact must be attenuated primarily in the lower extremities.

As a kind of soft tissues, muscles assist in the absorption of the impact force. Muscle activity can be tuned to impact force characteristics to control the soft-tissue vibrations^[8]^and protect the bone from external invasion ^**Error! Reference source not found**.^. Meanwhile, muscles are the major contributors to the joint moment acting at joints and on other parts of the body to contact with ground and objects to generate ^[10]^ the impact force ^[13]^and dynamic response.

Shock absorbing and cushioning properties of bone and soft tissues are also effective when a subject is not fatigued from impact loading ^**Error! Reference source not found**.^, however, bone fatigue or stress fracture is likely to occur under a higher impact loading during high-speed motion. Therefore, impact loadings generated during galloping has a higher influence on bone fatigue and stress fracture^[17]^. In present study, experimental method can be used to measure the stress-strain properties to obtain the dynamic response of bones^[20]^, other methods such as constructing the three dimensional models and numerical models of the musculoskeletal system by using special 3D capture system are also adopted^[21]^. However, there has been little work for dynamic modeling and experimental analyzing transient dynamic response of the lower extremities and the characteristics propagating along bones and soft tissues during the impact.

Therefore, in this paper, the musculoskeletal system model of a quadruped was firstly constructed from anatomical measurement of a domestic cat; Then Nano-indentation technology was used to measure the elastic modulus of the tibia precisely; Moreover, to obtain the transient response abilities of bones, soft tissues and bionic components, the substructure method was introduced in dynamics equations; Finally, impact experiments of tibia and a bionic leg were taken by high-speed videos system.

## Materials and Methods

### 1 Musculoskeletal system modeling

In this study, a domestic cat was obtained based on the laboratory animals welfare law. The domestic cat died of natural causes, and the body was kept under Zero degrees Celsius. In the anatomical data, the truck of the cat was 280 mm-long, shoulder was 60/70 mm-wide, the body weight was 1.4 kg, and other measured data were given in Table 1. Muscles were distributed along various joints and bones as shown inFig. 1a. Muscles act like linear motors to drive bones and joints, translating muscle contractions into rotations around bone joints. In the simplified model (Figure 1a), the attachment points of the tendons can be used to construct a kinematic model of the musculoskeletal system. Bones and joints are simplified according to their geometrical features and body motion functions.

**Fig. 1.**
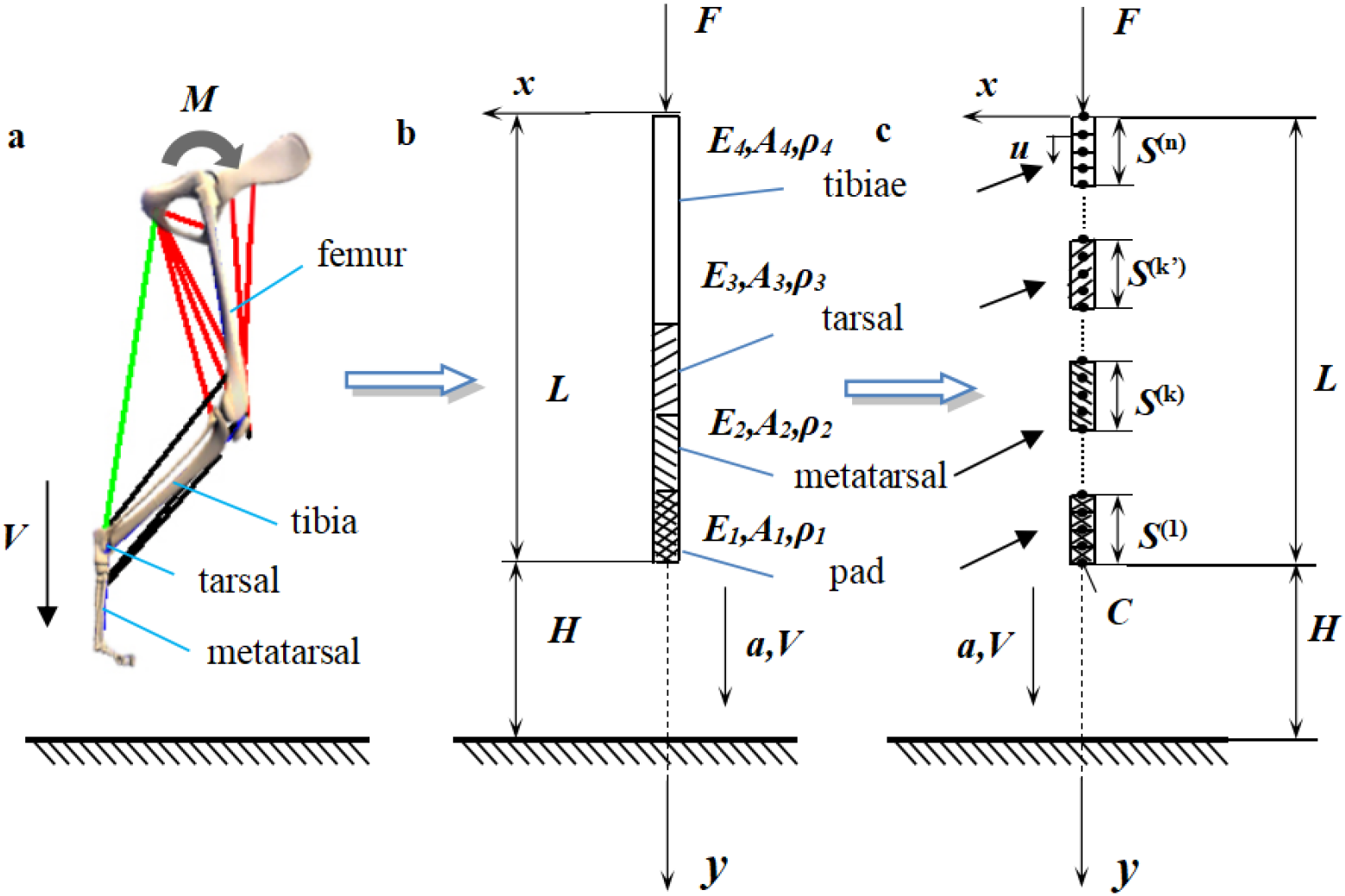
(a) The kinematic sketch of bionic mechanism musculoskeletal system and (b) the schematic figure of a tibia simpacting the ground and (c) the figure of the substructure used in the impact rod

To construct the kinematic model of the musculoskeletal system, the coordinate transformation method was employed:

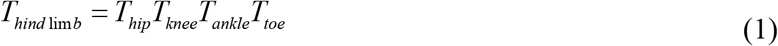

Eq. (1) was the kinematic equations of the hind limbs, and *T*_*hip*_, *T*_*knee*_, *T*_*ankle*_, *T*_*toe*_ are kinematic transform equations of the latter joint relative to the former joint. The kinematic equations of other limbs had the similar expression.

The muscle forces could be calculated by solving Eq. (2) as follows:

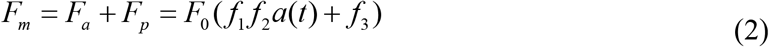

In Eq. (2), if the change of the distance between muscle tendon attachment points (Fig. 1b) varying with time is known, then Fm represents the size of muscle force represented by single connection lines shown in Fig. 1b could be obtained. In Eq. (2), ***F***_***a***_ and ***F***_***p***_ represent the forces of flexor muscle and extensor muscle respectively. ***f1*** 、 ***f2*** 、 ***f***_***3***_ represent the exponential function and is the arctangent function of the change in muscle length and contraction velocity respectively. The muscle length and the contraction velocity could be solved based on the kinematic model of the musculoskeletal system (Fig. 1a) and the measured running sequence of the cat.

Because of the different elastic moduli of the skeleton in Fig. 1a, the limb was simplified into the impact rod of different segments in Fig. 1b, the lower end of the rod represented the pad of the paw, the elastic modulus was *E*_*1*_, the segment next to the pad of the paw is the metatarsal bone, the elastic modulus is *E*_*2*_, then the next segment is the tarsal bone, the elastic modulus is *E*_*2*_, the third segment is the tarsal bone, the elastic modulus is *E*_*3*_, the remaining part is the tibia, the elastic modulus is *E*_*4*_. According to substructure method, this paper divided the impact rod into *n* substructure rods, each one containing *m* two nodes link element, as shown in Fig. 1c, *C* was a contact point, ***a***, ***V*** were the acceleration and speed of rod movement, *u* represents the physical displacement of node.

### 2 Determination of biomaterial properties

The tibia bridges the preceding and the following for the transmission of movement, where is a common site of stress fractures during running (Boutwell et al. 2015; Milgrom et al. 2000). Therefore, the tibia of the domestic cat is selected for this study (Fig. 2a).

**Fig. 2.**
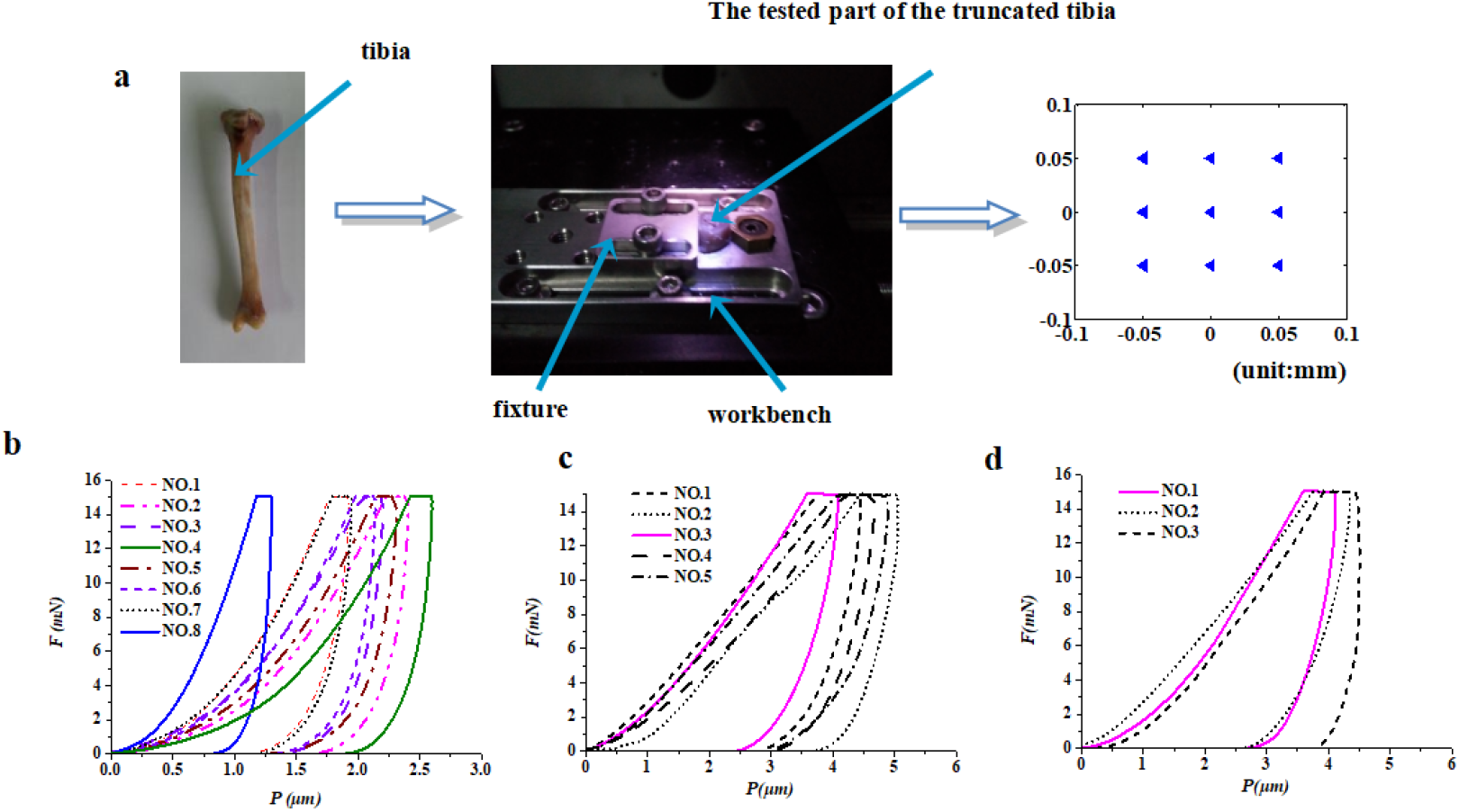
The nano-indentation experiment and the measured data (universal nano & micro tester, CETR, CA, USA). Fig. 2a shows the selected tibia, the workbench of the experiment, the tested points of the nano-indentation experiment. There are a total of 9 points in the tested area. The maximum pressed depth of the nano indentor is 5μm, light microscope resolution is 0.1μm, where the pressure accuracy is 0.1%, the mounted truncated tibia was also shown in Fig. 2a. Fig. 2b, c, d are the displacement-loading curves of different parts of tibia material; x axis was pressed depth, y axis was press-in force. Fig. 2b and Fig. 2c show the displacement-loading curves of tibia material, Fig. 2b is the nano-indentation data for the outside surface of tibia, Fig. 2c for the inside of tibia and medial condyle. There are eight groups of reasonable measured value in Fig. 2b, six in Fig. 2c. We fed the measuring results into the O&P method to obtain multi-group elastic moduli of tibial material, which were averaged to get the elastic moduli of the truncated tibia and the medial condyle as 7.45Gpa and 1.485Gpa respectively (Oliver etal. 1992). Fig. 2d showed the measured data of the medial malleolus (NO. 1 NO. 2) and tarsal (NO. 3), the elastic moduli were 1.429 Gpa and 2.579 Gpa respectively.

**Fig. 2.**
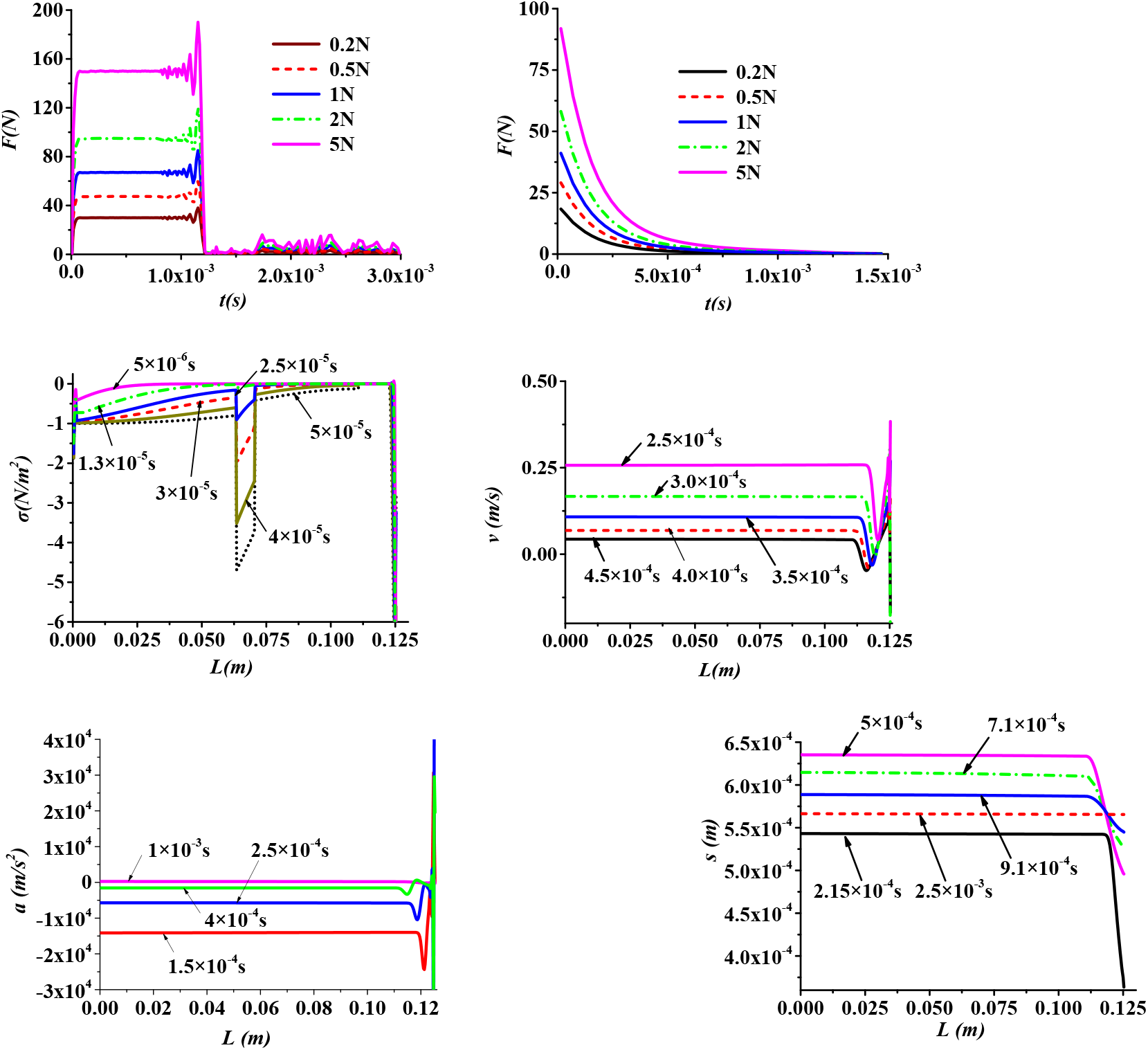
transient dynamic response of the lower limb (a), (b) the limb falling freely without and with considering the damping effects, (c) the stress wave distributed on the rod, (d), the velocity wave distributed on the rod, (e), the acceleration wave distributed on the rod, (f) the displacement wave.

The cat’s tibia is 83 mm long, the lateral ankle is 22 mm long, weight 1.4 g, with a density of about 791.6 kg/m^3^, the tarsal is 7.3 mm long, weight 0.52 g, with a density of about 1222.7 kg/m^3^, the metatarsal is 40 mm long, weight 2.41g, with a density of about 1652.7 kg/m^3^, the pad of the cat’s paw is 15 mm thick, with a density of about 1120 kg/ m^3^, the truncated tibia is 41 cm long, weight 1 g, with a density of about 1940.9 kg/m^3^, The experimental data were obtained after the tibia was soaked in physiological saline for more than 1 hour. The Young’s modulus were measured using a nano-indentation technology (Fig. 3) (universal nano & micro tester, CETR, CA, USA).

**Fig. 3.**
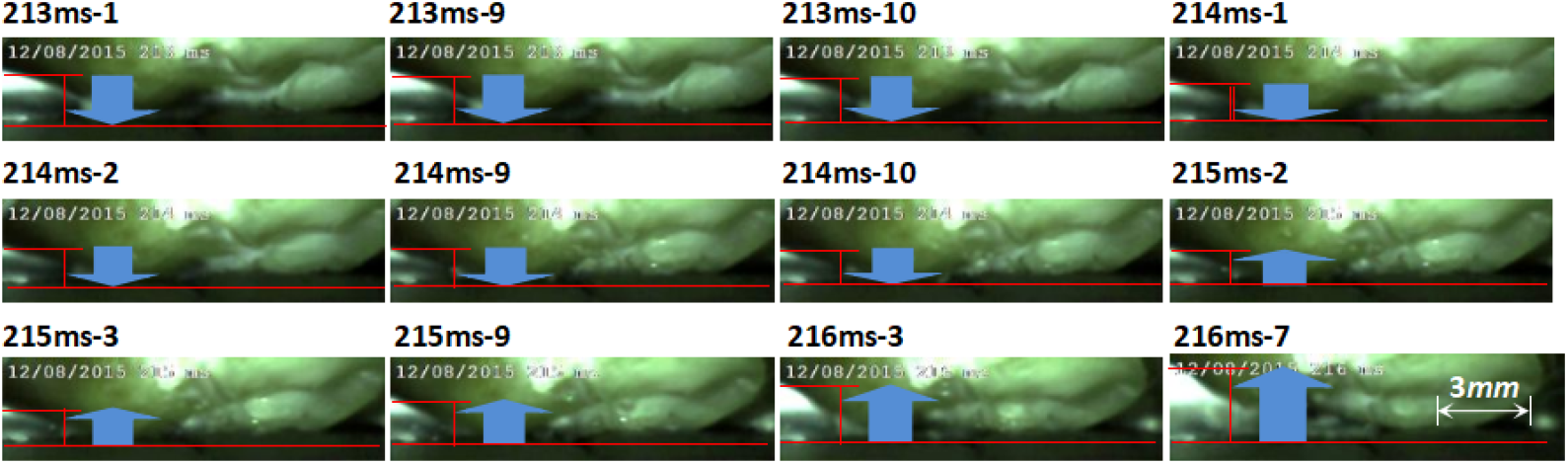
The tibia with soft tissues fell freely under a 100g load and impacted the rigid surface. From 213ms-1 and 214ms-10, the tibia fell down to impact the rigid surface. From 215 ms^-2^ and 216 ms^-7^, the tibia began to bounce off the rigid surface.

### 3 Numerical model of the dynamic response process

To investigate the transient response of bones, the contact between the tibia and tarsal was simplified: the tibia was simplified into one-dimensional plane rod, and the tarsal into rigid surface (Fig. 3a), under the action of the joint’s moment ***M***, the femur exerted pressure on the tibia, a brief contact-impact will occur between them to generate the dynamic response.

Because of the high nonlinear problem for solving dynamic response, the substructure technique for dynamics was employed, this method can reduce the modal order of finite element method, and can ensure the accuracy of the calculation (Fig. 3a). The dynamics equation of substructure ***S***^***(k)***^ can be expressed as:

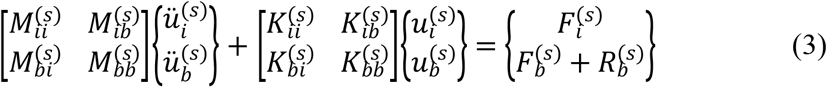

Namely:

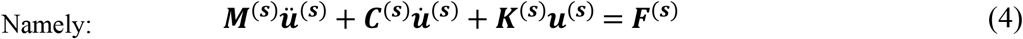

The mass matrix of Eq. (4) is a coordinated mass matrix, ***F*** is the external load, ***R*** is the interfacial force, and the physics displacement vector ***u*** is divided into two parts: the internal node displacement ***u***_***i***_^***(s)***^ and the interface node displacement ***u***_***b***_^***(s)***^. Through calculating the transforming relationship between physical displacement and modal displacement, the modal matrix could be obtained, then Eq. (4) could be solved based on the known parameter matrices. Meanwhile, during the rod vertically impacted the ground, it was assumed no invasion between the rod and ground, and no geometric dispersion along the rod. During the process of solution, the rod length *L* was divided into about 300 units, every four nodes for a substructure unit, and the convergence of the results of substructure method had been proved (Fig. 3c).

## Results

On the basis of above numerical model, the transient dynamic response properties of the lower limb of the cat was obtained (Fig. 2). The impact forces generated between the lower limb and the rigid surface were numerically calculated. During the impact, the lower limb was acted under five different external forces (Fig. 2a, b). It was obviously that with the increase of acting load, the impact force continued to increase. There was a damping effects during the lower limb impacted the rigid surface (Fig. 2b), the curves of the impact forces were declined rapidly and smoothly. When the object impacted the rigid surface without considering the damping effects, the impact forces would be varied with a rectangular waveform (Fig. 2a).

The stress wave produced by the impact could propagate through different bones and soft tissues of the lower limb. The tarsal endured the main pressure generated by the tibia and the metatarsal bone, where the stress value was obviously higher than that distributed on the tibia and the metatarsal bone. The velocity of the stress wave travelling through the paw pad was slower than that travelling through the skeletons (Fig. 2c). Therefore the velocity wave of the transient dynamic response travelling along the paw pad was slower. While, the amplitude change of the velocity wave was larger. The velocity wave distributed on the skeletons performed with a more identical variation, almost each point of the skeletons had the same velocity variation during the impact (Fig. 2d). The waveform of the acceleration was similar with velocity wave that distributed on the lower limb, the amplitude variation of which was significant at the paw pad (Fig. 2e).

Different from the propagation characteristics of velocity wave and acceleration wave, before reaching the upper end, the displacement wave distributed on the lower end of the limb, which travelled along the simplified rod with an almost linearly curve. At 2.15×10^−4^ s, the paw pad was kept contact with the rigid surface, and the upper end of the lower limb had been off the rigid surface with a displacement more than 5×10^−4^ m. During the impact, the displacement distributed on the bones varied with the similar trend, and at about 2.5×10^−3^ s, the lower limb had been ready to bounce off the rigid surface with an almost same displacement (Fig. 2f).

A high-speed video system (SHIQ-Motion-Pro-Y3, IDT, CA, USA) was used to record the dynamic response characteristics of the cat’s bones. The frequency of the high-speed video system is 10000 fps (frames per second), with 1:1 dedicated lens. With this frequency, in a millisecond (ms) numbered as u, there were 10 frames, which were labeled as u ms-1, u ms-2, …, u ms-10 (Fig. 3).

Under the effect of a 100 g load, an intact tibia with soft tissues fell freely and impacted the rigid surface (Fig. 3). During the impact, the soft tissues of the tibia continuous contacted the rigid surface. Under the action of the external forces, the tibia was firstly pressed, and then bounced off the rigid surface between 214ms-10 and 215ms-2. The displacement of the press process of the tibia was about 0.43 mm.

The transient dynamic response property of the impact ground was numerically solved. In the numerical simulation, two masses weighted 0.68kg and 0.34kg impacted the soil, the impacting velocity was 0.5 m/s and 0.25 m/s. The same mass falling down with different velocities would have the same variation periods of the impact forces, while the amplitudes were different (Fig. 4a). At 1186-ms, the mechanical leg (PLA material) was falling down with a certain load (Fig .4b). At 1200-ms, the mechanical leg began to bounce off. The mechanical leg stepped into the soft soil throughout the whole impact duration (from 1186-ms to 1211-ms).

**Fig. 4.**
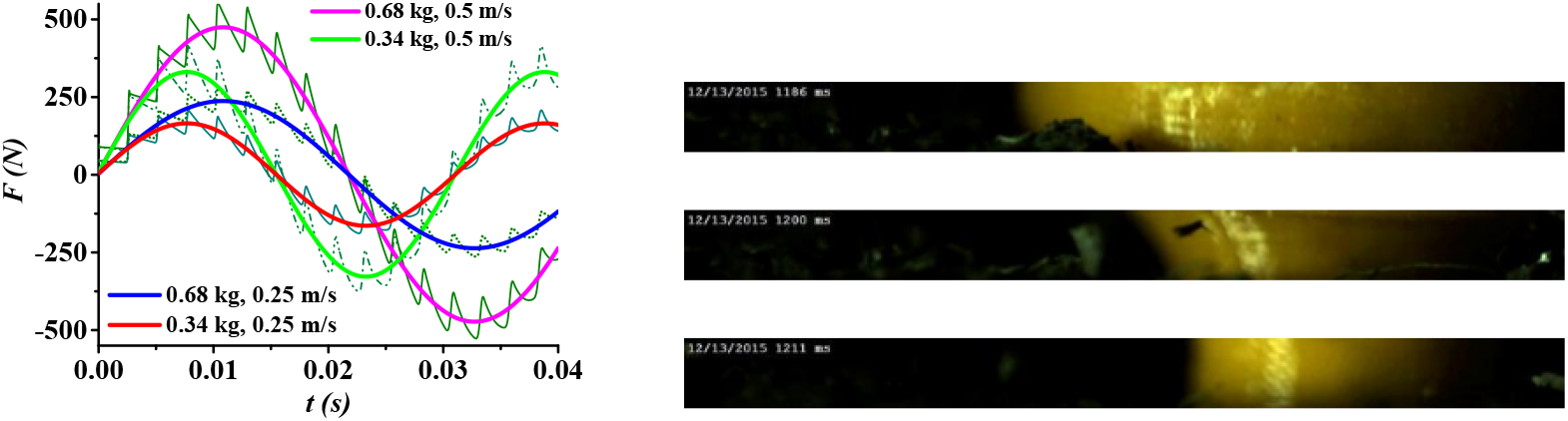
the transient dynamic response wave of the soil (a), the numerical results when the mass block impacting the soil, (b) the high speed video image of the leg stepping on the soil.

The small felid was trained to run through a line path and stepped on the strain gauge, the detected results showed that the duration of one step was about x ms during the fast running (velocity is about x m/s) (Fig. 6a). When the felid ran through the rigid ground, the impact duration of one step during the fast running was about y ms (Fig. 6).

## Discussions

As the primary motion part of the body for mammal, the musculoskeletal system is more sensitive to the stress response, so it was necessary to study the dynamic response characteristics of the primary skeleton of a quadruped. Therefore, the dynamic response of the lower limb of a domestic cat was studied.

The domestic cat was accurately measured. While, the skeleton consisted of cancellous bone and compact bone, the elastic modulus of cancellous bone was about one-third of that of compact bone in the measured data, and the cancellous bone had a larger contact area, in the connection area of joints. The skeleton of the lower limb had a good buffering and shock absorbing ability together with the soft tissues (such as the paw pad) ^[22]^.

The impact is a kind of transient dynamic response^[22]^, which is a high nonlinear problem that can be solved by using numerical or analytical method. In this study, the numerical method was used, the lower limb was simplified into a rod with or without different materials (e.g. soft tissues). The contact forces generated during the impact had a rectangular waveform varying with time (Fig. 2a)^[2]^. This is because the numerical model of the lower limb had a regular geometric body and uniform distribution of the material properties, not considering the damping effects ^[23]^. During the cat’s high-speed running, the soft tissues of the animal’s body can be considered as a vibrating system, and the damping effects can reduce the dynamic response during the vibration. Under considering the damping coefficient of the numerical model, the amplitude of the impact forces declined significantly (Fig. 2b)^[26]^. While the first impact duration was almost not influenced by the damping coefficient, it can be concluded that the frequency of the initial contact-impact would not vary much ^[28]^.

The transient response waves (stress wave, velocity wave etc) of the impact may have different distributed characteristics along the tibia. It is because that the material properties (e.g. ***E, ρ***) of the adjacent segments of the lower limb were different. When the stress wave propagated from the contact point C (Fig. 1c) to the upper end of the simplified rod, the stress wave will reflect back immediately at the upper end (Guo et al. 2004). The stress distributed on the tarsal was obviously larger than on the metatarsal and tibia. This is because the analytic solution of the amplitude of the impact force is: ***ρ***\**c*\**v*\*s, where***ρ***is the density of the rod, *v* is the initial impact velocity, *s* is the contact area. In the numerical model of this study, the density and moduli of different bones had little differences, while the contact area of the tarsal was more larger than the adjacent bones (Escalona et al. 1999). However, in the cat’s lower limb, the contact face between two adjacent bones are connected as a joint, where the contact faces of the tarsal were curved surfaces, and the tarsal were almost embedded in the ankle joint of the cat, therefore the tarsal could endure greater impact stress.

Compared with the stress wave, the change of the displacement wave was relatively smaller, especially when the displacement wave travelled across the interface of different bones. The displacement distributions along the rod at different time instants during the impact were numerically solved (Fig. 2f). The stress wave originated at instant and the rod deformed at the contact surface. With time progressing, the stress waves were generated and propagated along the rod at different velocity in different sections of the lower limb (Fig. 2d). The displacement and the deformation of the section on the rod that the waves had already passed increased correspondingly, especially the deformation of the section of soft tissues changed significantly (Fig. 2f) and the left sections showed almost no significant deformations. However, the displacement of the lower limb had always been changing during the impact^[2]^. It could be concluded that the deformation of each segment of the lower limb had not been achieved to the yield strain, which included the yield strain of the soft tissues, cartilage, and bones^[21]^. Namely, the lower limb is in the elastic deformation state, which would benefit the high speed running ability of the cat. The velocity wave, acceleration wave had the similar distribution on the lower limb, where the variations in the soft tissues was larger than in other segments (Fig. 2d, e).

The tibia with some soft tissues was selected to impact the rigid surface under a certain load (Fig. 3). The initial impact took about 2-ms, which is obviously longer than the solved results. This is because that the elastic modulus distributed on the tibia were stable and non-uniform, while the elastic modulus and densities of the cancellous bone, compact bone were different. The cartilage and soft tissues (such as muscle, paw pad) all had a good shock absorbing and buffering ability. Therefore, the lower limb with different joints and skeletons of the cat could have a longer impact duration than the numerical results.

From the numerical results, it can be seen that the impact duration (about 20ms) between the mechanical leg and the soil (Fig. 4a) was similar with the results calculated (0.68kg, near 20ms, the first impact) (Fig. 2b). At the end of the falling down period, the mechanical leg almost did not keep contact with the soft ground, and bounced off immediately. It could be concluded that there was an energy feeding back process. Namely, the soft soil could provide elastic potential energy for the limb during the high speed running ^[28]^. Based on this view point, the elastic deformation of the lower limb of the cat could also provide the energy for its high speed running. This is why one should expect resonant oscillations to occur during the running ^**Error! Reference source not found**.^.

The tested duration of the felid’s one step during the fast running were similar with the response duration of the soil, it suggests that the impact duration is determined by the material with a longer response duration, though the conclusion need to be further investigated. When the felid runs through the rigid ground, the duration of the one step during the fast running was close to the results of the first impact duration of the lower limb. It suggests that the felid’s running frequency was similar with the computed results that generated between the lower limb and the rigid surface. Therefore the resonant may happen to the lower limb during the felid’s high speed running^[25]^.

## Conclusions

The transient dynamic response of the felid’s lower limb was numerically solved and experimentally verified during the impact. The initial impact forces were relatively higher, which was only related to the mechanical property of the lower limb (E, ρ, etc.). The impacts exist between limbs and the soil may provide the feedback energy for the felid’s agile movement. The properties of the transient dynamic response between the limbs and the ground (soil) can be employed to improve further analyzing the function of the musculoskeletal system during the movement.

## Acknowledgement

This study was supported by the Tianjin Education Commission Research Project (2022KJ124).

